# Is shape in the eye of the beholder? Assessing landmarking error in geometric morphometric analyses on live fish

**DOI:** 10.1101/2022.10.11.509915

**Authors:** Paolo Moccetti, Jessica R. Rodger, Jonathan D. Bolland, Phoebe Kaiser-Wilks, Rowan Smith, Andy D. Nunn, Colin E. Adams, Jen A. Bright, Hannele M. Honkanen, Angus J. Lothian, Matthew Newton, Domino A. Joyce

**Affiliations:** Evolutionary and Ecological Genomics Group, School of Natural Sciences, University of Hull, UK; Hull International Fisheries Institute, School of Natural Sciences, University of Hull, UK; Energy and Environment Institute, University of Hull, UK; Atlantic Salmon Trust Fellow, Scottish Centre for Ecology and the Natural Environment, Institute of Biodiversity, Animal Health and Comparative Medicine, University of Glasgow, UK; Scottish Centre for Ecology and the Natural Environment, Institute of Biodiversity, Animal Health and Comparative Medicine, University of Glasgow, UK; School of Natural Sciences, University of Hull, UK

**Author notes:** Corresponding Author: Paolo Moccetti^1,2,3^, School of Natural Sciences, University of Hull Cottingham Road, Hull, HU6 7RX, UK, Email address.

## Abstract

Geometric morphometrics is widely used to quantify morphological variation between biological specimens, but the fundamental influence of operator bias on data reproducibility is rarely considered, particularly in studies using photographs of live animals taken under field conditions. We examined this using four independent operators that applied an identical landmarking scheme to replicate photographs of 291 live Atlantic salmon (*Salmo salar* L.) from two rivers. Using repeated measures tests, we found significant inter-operator differences in mean body shape, suggesting that the operators introduced a systematic error despite following the same landmarking scheme. No significant differences were detected when the landmarking process was repeated by the same operator on a random subset of photographs. Importantly, in spite of significant operator bias, small but statistically significant morphological differences between fish from the two rivers were found consistently by all operators. Pairwise tests of angles of vectors of shape change showed that these between-river differences in body shape were analogous across operator datasets, suggesting a general reproducibility of findings obtained by geometric morphometric studies. In contrast, merging landmark data when fish from each river are digitised by different operators had a significant impact on downstream analyses, highlighting an intrinsic risk of bias. Overall, we show that, even when significant inter-operator error is introduced during digitisation, following an identical landmarking scheme can identify morphological differences between populations. This study indicates that operators digitising at least a sub-set of all data groups of interest may be an effective way of mitigating inter-operator error and potentially enabling data sharing.

## Introduction

Landmark-based geometric morphometrics (GM) is a quantitative approach widely used to describe the shape of biological specimens and its covariation with other biological and environmental factors (Zelditch et al., 2004; Webster & Sheets, 2010). Morphological variables are quantified using a set of Cartesian landmarks located on distinct homologous anatomical points, and observed body shape variations are then displayed through user-friendly graphical representations (Zelditch et al., 2004; Mitteroecker & Gunz, 2009; Adams, Rohlf & Slice, 2013). GM is a powerful technique capable of detecting even tiny morphological differences among groups of specimens (Mitteroecker & Gunz, 2009; Webster & Sheets, 2010), but is highly sensitive to measurement errors introduced during data acquisition, which can affect subsequent analyses and produce inaccurate results (von Cramon-Taubadel, Frazier & Lahr, 2007; Fruciano, 2016; Robinson & Terhune, 2017; Fox, Veneracion & Blois, 2020). This is particularly problematic when such morphological differences are erroneously regarded as biologically meaningful variation (Fruciano, 2016).

Surprisingly, despite GM being a widely used technique, researchers rarely consider measurement error in their study design and statistical analyses (Fruciano, 2016; Fox, Veneracion & Blois, 2020). Measurement error can be introduced at different stages of the data acquisition process, i.e. when positioning specimens in front of the imaging device (camera or scanner), during image capture and landmark digitisation (Arnqvist & Mårtensson, 1998; Muir, Vecsei & Krueger, 2012; Fruciano et al., 2020; Fox, Veneracion & Blois, 2020). Indeed, the so-called inter-operator (or inter-observer) error during landmarking has been found to be one of the most critical factors affecting GM analyses because different operators tend to position what should be homologous landmarks in slightly different locations (Ross & Williams, 2008; Dujardin, Kaba & Henry, 2010; Campomanes-Álvarez et al., 2015; Fruciano, 2016; Fruciano et al., 2020; Fox, Veneracion & Blois, 2020).

Importantly, inter-operator error can be substantial and potentially obscure biological variation, making data sharing and comparisons of landmarked datasets difficult (Shearer et al., 2017). Intra-operator (or intra-observer) error has also been shown to significantly affect GM analyses (Wilson, Cardoso & Humphrey, 2011; Fox, Veneracion & Blois, 2020). Intra-operator error is introduced when specimens are inconsistently digitised by a single operator and can be influenced by several factors, including landmarking experience or time between landmarking sessions (Fox, Veneracion & Blois, 2020). However, the magnitude of intra-operator error is invariably modest compared to inter-operator discrepancies (Cardoso & Saunders, 2008; Dujardin, Kaba & Henry, 2010; Wilson, Cardoso & Humphrey, 2011; Robinson & Terhune, 2017; Shearer et al., 2017; Thoma et al., 2018; Fox, Veneracion & Blois, 2020), indicating a general good precision in digitisation by individual operators (but see Engelkes et al., 2019).

The degree and impacts of operator error in GM studies have been tested for a range of organisms, anatomical structures, preservation methods and image acquisition devices (Fruciano, 2016; Fruciano et al., 2020; Fox, Veneracion & Blois, 2020). Nevertheless, most studies have focussed on images of specific human, bone or plant structures acquired under identical (laboratory) conditions (e.g., Ross & Williams, 2008; Cardoso & Saunders, 2008; Gonzalez, Bernal & Perez, 2011; Wilson, Cardoso & Humphrey, 2011; Viscosi & Cardini, 2011; Shearer et al., 2017; Carayon et al., 2019; Engelkes et al., 2019; Messer et al., 2021). Few have investigated images of live animals (but see Fruciano et al., 2020), despite commonly being used when it is not possible to euthanise samples for ethical reasons or research purposes. Undeniably, such photographs, especially if taken under field conditions, are more likely to result in subsequent measurement error (relative to preserved specimens) (Muir, Vecsei & Krueger, 2012), thereby restricting the utility of such datasets (Webster & Sheets, 2010). Understanding the prevalence, magnitude and implications of inter- and intra-operator error during the landmark digitisation process for photographs of live animals could facilitate data sharing and open science practices.

With the increasing focus on reproducibility in science (Baker, 2016), and an acknowledgment that sharing data can accelerate scientific progress, assessing whether live animals digitised repeatedly by single versus multiple operators produce consistent results and conclusions is essential. Data exchange, such as crowdsourcing, is opening new frontiers in GM research, enabling large-scale studies, which use unprecedented sample sizes, to be conducted within a short time frame (Thomas, Bright & Cooney, 2016; Chang & Alfaro, 2016). Such studies, involving several operators collecting shape data, can potentially address key questions in evolutionary biology and other disciplines (Cooney et al., 2017; Hughes et al., 2022). However, pooling landmarked datasets from multiple operators can increase the degree of measurement error (Fruciano et al., 2017; Evin, Bonhomme & Claude, 2020), but the consequences of inter-individual operator error when sharing datasets remain poorly understood.

The aim of this study was therefore to determine whether GM analyses on photographs of live animals are reproducible. To accomplish this, four independent operators digitised the same photographs of sedated Atlantic salmon (*Salmo salar* L.) sampled in two rivers, following a shared landmarking scheme. The shape data and results obtained by the four operators were then compared and contrasted to assess the magnitude of inter- and intra-operator error, and infer the potential for meaningful data sharing.

## Material & Methods

### Study design

Salmon were captured from the River Spey (57° 24.960’ N 3° 22.602’ W) and River Oykel (57° 59.640’ N 4° 48.282’ W) in Scotland using a 1.5 m diameter Rotary Screw trap during their smolt stage, i.e. on their first migration to sea. The sampling occurred in the context of a tracking study aiming to identify areas and causes of smolt mortality during their seaward migration (see Whelan, Roberts & Gray, 2019; and https://atlanticsalmontrust.org/our-work/morayfirthtrackingproject/). Fish were photographed in the field under anaesthetic before being tagged and released to the river after recovery. Photographs of the left side of each fish were taken freehand from approximately 30 cm directly above the fish, with a Fujifilm FinePix XP130 Compact Digital Camera with fish on a background reference scale. Photographs were taken by a team of eight people who met prior to field work to standardise methods as far as possible. Our study here focusses on inter-operator variation downstream of photography, but variation caused by variation between individual photographers would be worthy of future study. The care and use of experimental animals complied with the UK Home Office animal welfare laws, guidelines and policies (UK Home Office Licence PPL 70/8794) and was approved by the University of Glasgow Animal Welfare and Ethics Review Board (AWERB). Field permits were provided verbally by: Keith Williams: Kyle of Sutherland Fisheries Trust (River Oykel) and Brian Shaw: Spey District Salmon Fisheries Board (River Spey). The GM analyses were based on photographs of 291 salmon (Spey *n* = 144, Oykel *n* = 147). The images were imported into tpsUtil v. 1.78 (Rohlf, 2019) and randomly shuffled using the relevant function so that operators were blinded to the river-of-origin of the specimens.

Fifteen fixed and seven semi-landmarks (Bookstein, 1997) were digitised on each image by four independent operators (Op.1, Op.2, Op.3 and Op.4) using tpsDig v. 2.31 (Rohlf, 2017) and following an identical scheme (Fig. 1). The landmark positions chosen were those commonly used in studies on salmonids (e.g., Boulding et al., 2008; Muir, Vecsei & Krueger, 2012; Simonsen et al., 2017; Goerig et al., 2019; Dermond, Sperlich & Brodersen, 2019). In addition, the first ten fish for each river, after using the *randomly order specimens* function in tpsUtil, were consecutively landmarked a further two times (i.e. three times in total) by each operator to evaluate the intra-operator consistency in digitisation.

**Figure 1.**
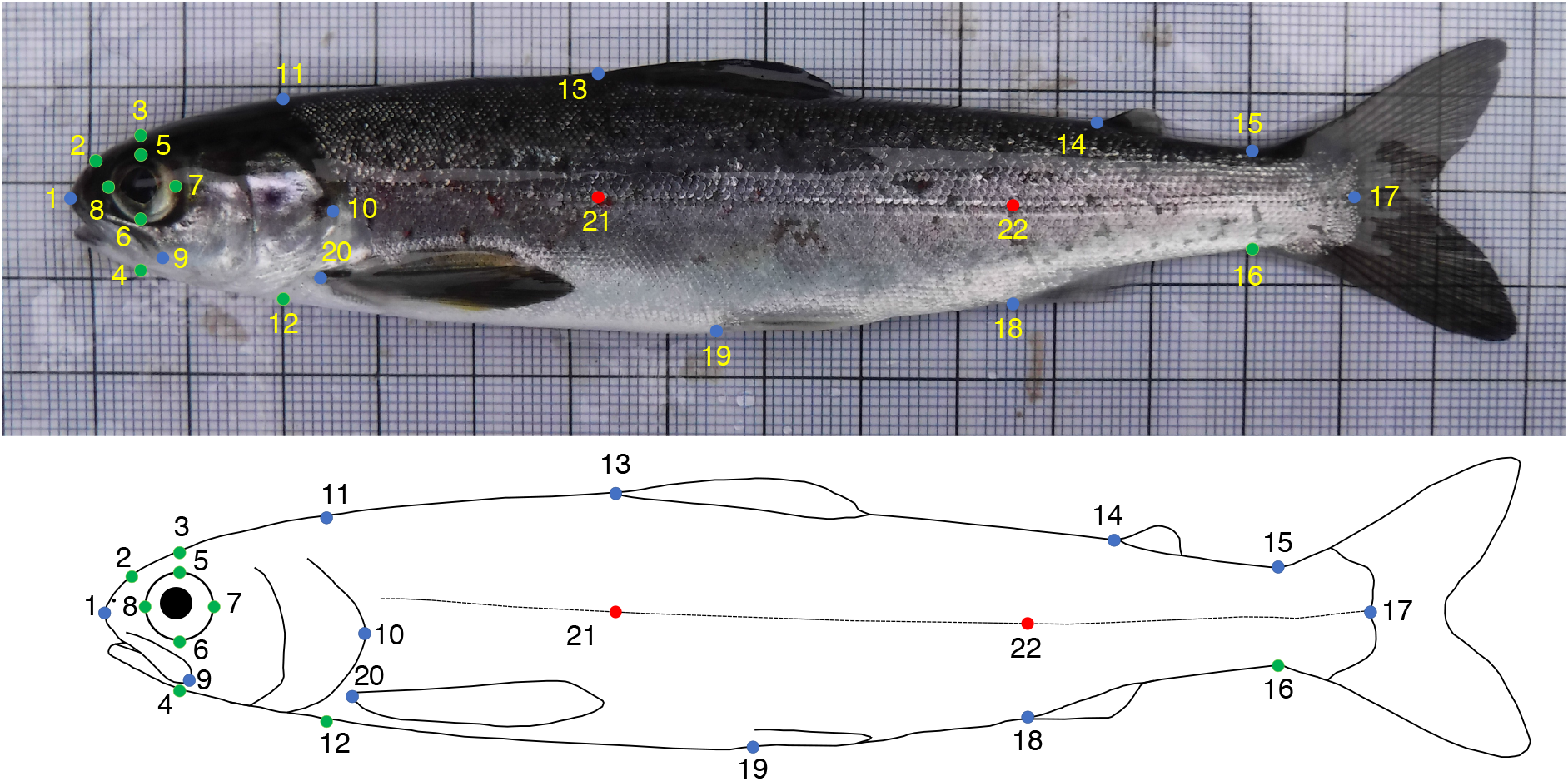
Fixed (blue) and semi-(green) landmarks used for the geometric morphometrics analyses of Atlantic salmon smolts. Landmarks 21 and 22 (red) were used to correct for body arching and not included in the analyses. (1) Tip of snout; (2) Midpoint between 1 and 3; (3) Directly above middle of eye; (4) Perpendicular to 3; (5) Midpoint of top of eye (directly below 3); (6) Midpoint of bottom of eye (directly below 3); (7) Midpoint of posterior of eye; (8) Midpoint of anterior of eye; (9) End of maxillary bone; (10) Posterior tip of bony operculum; (11) Dorsal surface posterior of cranium; (12) Perpendicular to 11; (13) Anterior insertion point of dorsal fin; (14) Anterior insertion point of adipose fin; (15) Dorsal insertion point of caudal fin; (16) Versal insertion point of caudal fin; (17) Posterior midpoint of hypural plate; (18) Anterior insertion point of anal fin; (19) Anterior insertion point of ventral fin; (20) Anterior insertion point of pectoral fin; (21) Lateral line - perpendicular to 13; (22) Lateral line - perpendicular to 18. Anterior insertion point of ventral fin; (20) Anterior insertion point of pectoral fin; (21) Lateral line - perpendicular to 13; (22) Lateral line - perpendicular to 18.

Landmark coordinates from all operators were imported as unique files into R (R Core Team, 2021) and analysed using the ‘geomorph’ and ‘RRPP’ v. 4.0.4 (Adams et al., 2021; Baken et al., 2021; Collyer & Adams, 2021), ‘Morpho’ v. 2.8 (Schlager, 2017), and ‘GeometricMorphometricsMix’ v. 0.0.8.4 (Fruciano, 2018) packages. Plots were produced with the ‘ggplot2’ package (Wickham, 2016), while projections of body shape variation between groups were generated with the *plotRefToTarget* function in ‘geomorph’.

The landmark data were then used to test if: (1) similar mean body shapes were obtained by all operators; (2) any morphological differences between salmon from the two different rivers were detected by all operators; (3) identified between-river differences were consistent across all operators; (4) divergent datasets from different operators could be merged; and (5) the magnitude of intra-operator error was similar across operators.

### Preliminary analyses

First, a generalised Procrustes analysis (GPA) with sliding of semi-landmarks was performed to remove effects not related to body shape through translation, scaling and rotation of the landmark configurations (Rohlf & Slice, 1990). A preliminary principal component analysis (PCA) conducted on superimposed coordinates revealed body bending as a major source of shape variation, a known issue in morphometric studies on fish (Valentin et al., 2008). To remove the bending effect, the *unbend* function in tpsUtil was used, employing landmarks 1, 21, 22 and 17, which normally lie in a straight line in salmonids (Arbour, Hardie & Hutchings, 2011; Dermond, Sperlich & Brodersen, 2019). All subsequent analyses were performed on landmarks 1-20 only. A new GPA on coordinates with the bending deformation removed was then executed and outlier specimens were investigated for each operator using the *plotOutliers* function in ‘geomorph’. Two specimens digitised by one operator were found to be very different to the other individuals and were therefore removed from the dataset of all four operators, leaving 289 samples for analyses (Spey *n* = 144; Oykel *n* = 145). Another GPA using the landmark data without outliers was then implemented.

### Test 1. Were similar mean body shapes obtained by all operators?

To investigate whether results produced by a single operator are accurate and reproducible, we tested differences in the mean body shapes of fish digitised by independent operators. First, a between-group PCA (Boulesteix, 2004) was computed to explore variations between the four operators. Between-group PCA is a type of discriminant analysis used to maximise segregation between known groups which, unlike canonical variate analysis (CVA), does not overestimate the degree of distinction among groups (Mitteroecker & Bookstein, 2011). The leave-one-out cross-validation operation was implemented to quantify the proportion of fish specimens correctly assigned to the operator who digitised them.

To investigate whether landmarking by multiple operators introduced bias, i.e. systematic error affecting body shape (*sensu* Fruciano, 2016), differences in the mean body shapes of the fish digitised by the four independent operators were tested using Hotelling’s *T*^2^ as implemented by the *repeated_measures_test* function in ‘GeometricMorphometricsMix’. To compute the differences in mean body shapes, a PCA was performed on all Procrustes-aligned coordinates of all operators, and the scores for all the PC axes (i.e. 100% variance explained) of each operator were then used in a repeated measures test as an approximation of shape.

### Test 2. Were morphological differences between salmon from different rivers detected by all operators?

We next tested whether there was a difference in body shape between rivers, and whether the operators were consistent in identifying any differences. The following analyses were performed separately for each operator. First, a GPA was computed on landmark coordinate datasets obtained by each operator with outliers removed (see end of *Preliminary analyses*). The effect of fish size on body shape was tested using Procrustes ANOVAs (*procD*.*lm* function in ‘geomorph’), with Procrustes coordinates used as an outcome variable, the log value of centroid size and ‘River’ as independent variables and a randomised residual permutation procedure (10,000 iterations). A small but significant effect of size on shape was found for all operators (*P*-values < 0.0001, *r*^2^ = 0.022–0.034). Procrustes coordinates were therefore adjusted for allometry by using residuals from a regression of shape against centroid size + ‘River’. Procrustes ANOVAs were then used to compare mean body shape between rivers, while another between-group PCA was implemented to quantify the proportion of fish correctly assigned to the river of origin, for each of the four datasets.

### Test 3. Were identified between-river differences consistent across all operators?

To assess if body shape differences between rivers were analogous across operators, pairwise angles (Li, 2011) of vectors of shape change between fish from the rivers Spey and Oykel were computed. The *TestOfAngle* function in ‘GeometricMorphometricsMix’ based on the analogous function implemented in ‘Morpho’ was used, as performed by Fruciano et al. (2020). Specifically, we calculated the pairwise angles among between-group principal components obtained using ‘River’ as the grouping factor within each operator subset of digitisations (one between-group PC axis - herein bwgPC - per operator) to test if they followed the same “direction”, i.e. if the shape differences between rivers were approximately the same for all operators.

Furthermore, bwgPC1 vectors of between-river differences for each operator were compared (test of angles) with vectors of inter-operator differences obtained by subtracting corresponding Procrustes corrected coordinates of each specimen (eg. [coordinates of specimen 1 digitised by Op.1] – [coordinates of specimen 1 digitised by Op.2]) and then calculating mean shapes for all specimens. In this way, it was possible to determine whether or not biological body shape differences between rivers and artefactual variation among operators were similar (following the same “direction”).

Finally, the magnitude of shape differences between rivers obtained by each operator was examined with the *dist_mean_boot* function in ‘GeometricMorphometricsMix’. This function was used to perform a bootstrap estimate of the shape distance between the two rivers and allowed us to test if the amount of shape difference between the rivers Spey and Oykel was consistent across different operators or, on the contrary, one or more operators detected larger or smaller between-river differences than the others.

### Test 4. Can divergent datasets from different operators be merged?

The two operators producing the most dissimilar mean shapes were used to simulate a worst-case-scenario process of data pooling, in which two independent researchers perform their own GM study each on different rivers, but following the same landmarking scheme. Inter-operator analysis showed that Op.2 and Op.4 produced the most dissimilar body shapes (greatest Euclidean distance), so from these, two datasets were created: one comprising shape data from the River Oykel digitised by Op.2 (herein Op.2-Oykel) and the River Spey data digitised by Op.4 (herein Op.4-Spey) and *vice versa*, i.e. the River Spey data digitised by Op.2 (herein Op.2-Spey) and the River Oykel data digitised by Op.4 (herein Op.4-Oykel).

For both datasets, differences between rivers were tested with Procrustes ANOVA, as described earlier. Then, a between-group PCA was performed and the resulting bwgPC1 separating the two rivers was used to run a test of angles to compare between-river differences detected by these two separate datasets. We also compared these latter between-river differences with those found when using the complete datasets of all four operators including both rivers (see section above). This enabled us to test whether any between-river differences as a result of different operators outweighed any biological differences between rivers found when using the complete intra-operator datasets.

### Test 5. Quantifying intra-operator error

A GPA was computed separately on landmark coordinates obtained by each operator re-digitising a sub-sample of 20 fish (ten per river). Individual consistency in landmarking was then investigated using PCA and tested using repeated measures tests. To test for differences in mean body shapes between digitisation trials, a PCA was performed on the Procrustes-aligned coordinates of each operator separately and the PC scores of each trial were then used in the repeated measures tests as an approximation of shape. Repeatability among digitisation trials was also calculated for each operator using the intraclass correlation coefficient (Fisher, 1958). A one-way Procrustes ANOVA was computed using individual fish as a categorical variable (Fruciano, 2016). The resulting mean squares were used to calculate repeatability by applying equations presented in Arnqvist & Mårtensson (1998) and Fruciano (2016). Here, repeatability measured variation in the three independent digitisations of the sub-sample of 20 salmon relative to the variability among specimens, i.e. the biological variation among all fish samples. Repeatability assumes a value of between zero and one, with one indicating 100% repeatability and an absence of measurement error (Arnqvist & Mårtensson, 1998; Fruciano, 2016). Finally, a Procrustes ANOVA with individual fish specimens (‘ID’) as the main factor and ‘operator’ nested within ‘ID’ was run to test the relative contributions of biological variation (‘ID’) and variation introduced by inter-operator (‘ID:operator’) and intra-operator (residual) error.

## Results

### Test 1. Were similar mean body shapes obtained by all operators?

Despite digitising replicate photographs with homologous landmarks, fish specimens were correctly assigned to their operator based on body shape with 83.0% accuracy by the exploratory between-group PCA (Fig. 2, Supplementary Table 1). There was a significant operator effect on mean body shape, with all pairwise tests displaying highly significant differences between operators (*P*-value < 0.001 for all comparisons; Table 1), supporting the exploratory between-group PCA (Fig. 2). The Euclidean distances between means, i.e. the measure of the extent of shape change, highlighted different distances among pairs of operators, with the smallest difference (0.00771) occurring between Op.1 and Op.3 and the greatest between Op.2 and Op.4 (0.01577). The between-group PCA scatterplot (Fig. 2) broadly reflected these results along axis 1 (61.9% of variance), with Op.1 and Op.3 overlapping extensively and Op.2 and Op.4 being furthest apart. The anatomical differences among operators were concentrated mainly on the head (Fig. 3, Supplementary Fig. 1), with major areas of disagreement being the snout, eye, mouth and posterior of the cranium.

**Table 1.** Pairwise comparisons of the body shape of 289 Atlantic salmon landmarked by four independent operators based on Hotelling’s *T*^2^.

| Comparison | Euclidean dist. | Hotelling's $T^2$ | $F$ | $P$-value |
| --- | --- | --- | --- | --- |
| Op.1 vs. Op.2 | 0.01023 | 7100.15 | 162.84 | $< 1 \times 10^{-6}$ |
| Op.1 vs. Op.3 | 0.00771 | 3422.47 | 78.49 | $< 1 \times 10^{-6}$ |
| Op.1 vs. Op.4 | 0.01135 | 5130.73 | 117.67 | $< 1 \times 10^{-6}$ |
| Op.2 vs. Op.3 | 0.01215 | 9760.50 | 223.86 | $< 1 \times 10^{-6}$ |
| Op.2 vs. Op.4 | 0.01577 | 7511.23 | 172.27 | $< 1 \times 10^{-6}$ |
| Op.3 vs. Op.4 | 0.01114 | 5910.24 | 135.55 | $< 1 \times 10^{-6}$ |

**Figure 2.**
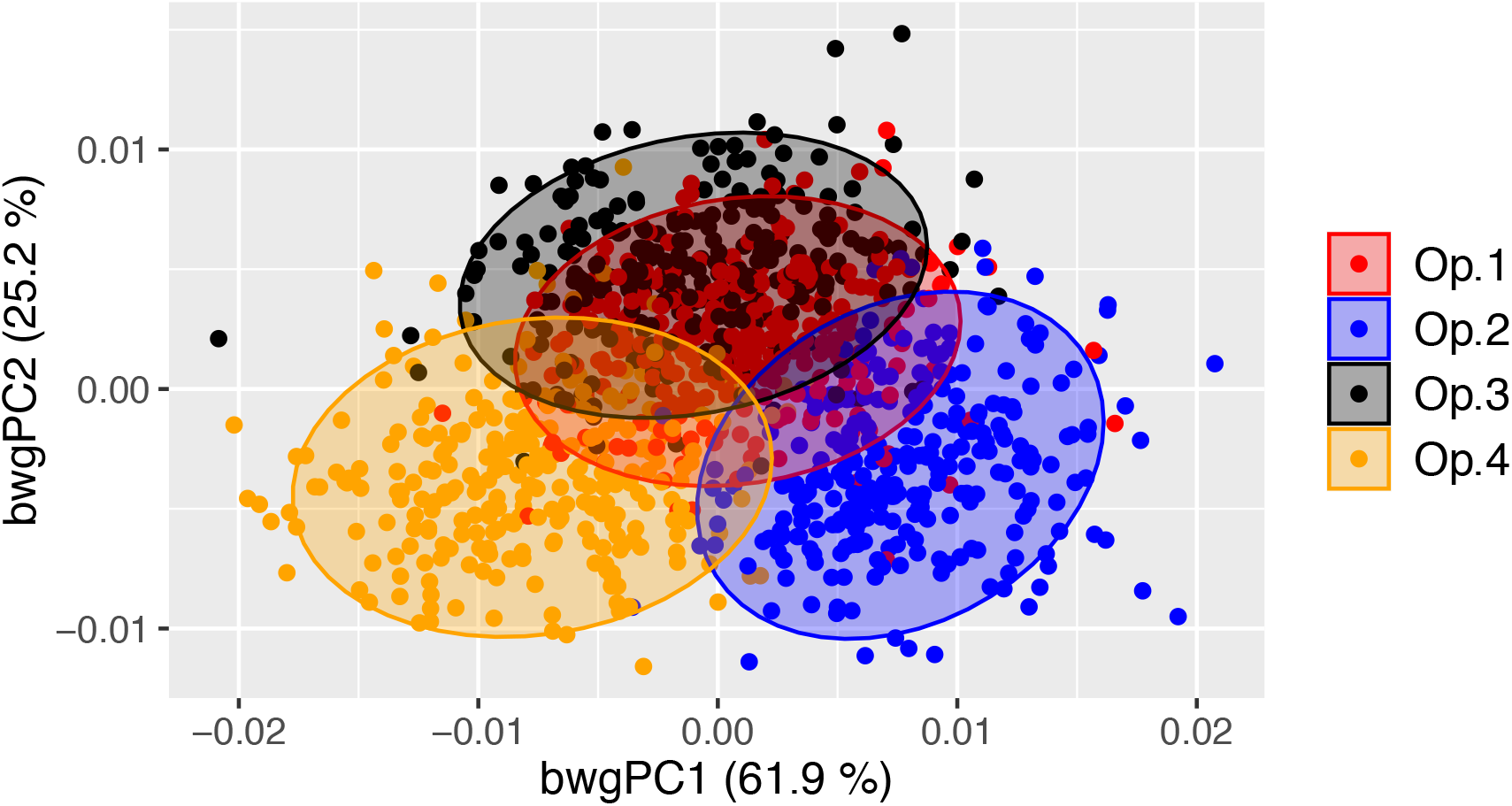
Between-operator PCA scatterplot showing the cross-validated scores along the first two between-group principal components (bwgPCs). Dots represent individual Atlantic salmon (*n* = 289) landmarked by four independent operators (different colours). Between-operator variance (%) explained by the first and second axes is shown.

**Figure 3.**
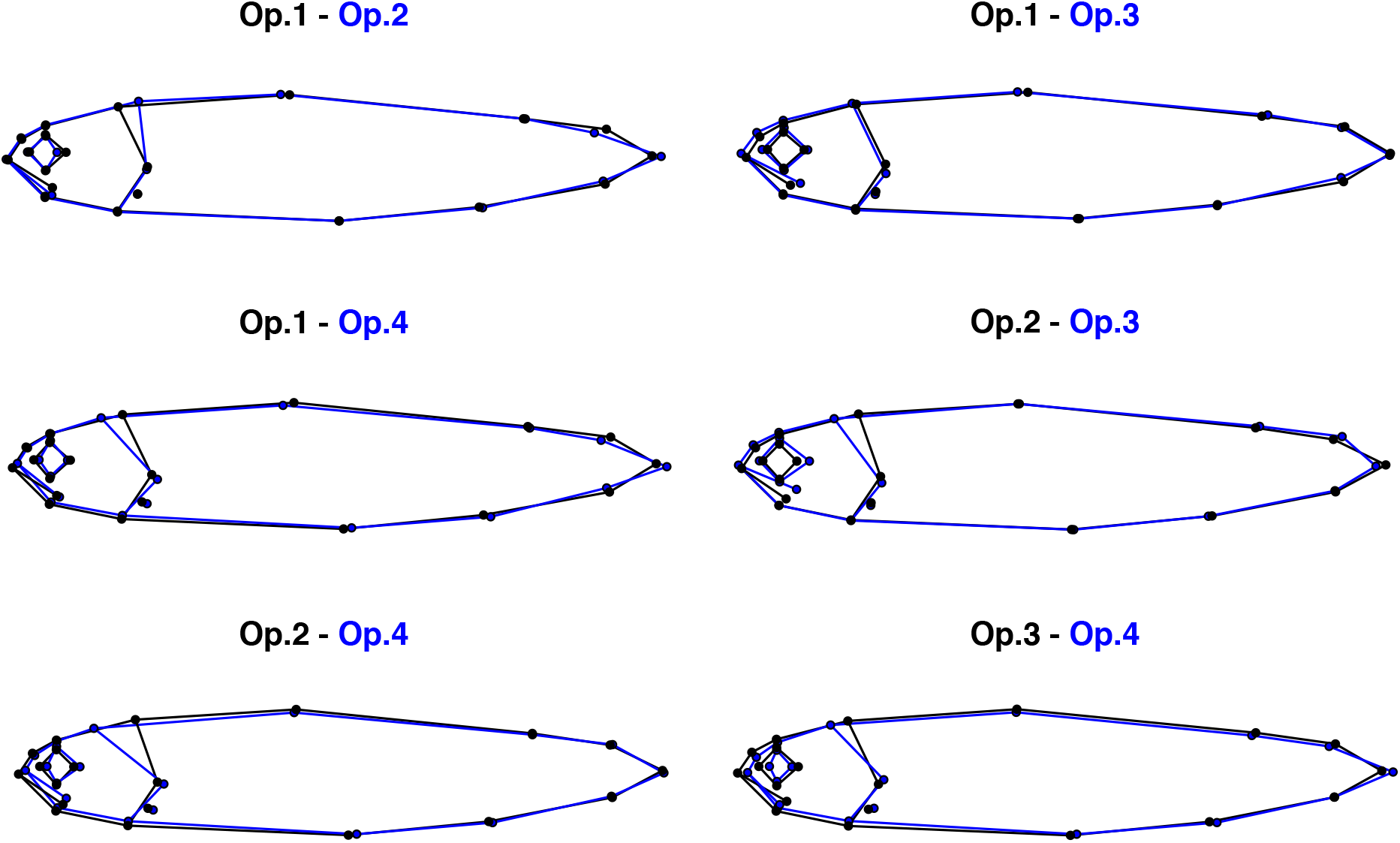
Pairwise comparisons of the mean body shape of 289 Atlantic salmon landmarked by four independent operators. Morphological differences were magnified five times to aid visualisation.

Morphological dissimilarities were more or less pronounced depending on the operator comparisons (Fig. 3).

### Test 2. Were morphological differences between salmon from different rivers detected by all operators?

There were significant differences in body shape between fish from different rivers (Spey and Oykel; Table 2), with between-group PCA (Supplementary Table 2) separating them for all operators (70.9% mean classification success rate). The fish from the River Oykel had a greater body depth, more pronounced caudal peduncle, larger eye, longer mouth and more pointed snout than those from the River Spey (Fig. 4).

**Table 2.** Procrustes ANOVA summary statistics of effect of river-of-origin on the body shape of 289 Atlantic salmon landmarked by four independent operators.

| Operator | Df | SS | $r^2$ | $F$ | $Z$ | $P$-value |
| --- | --- | --- | --- | --- | --- | --- |
| Op.1 | 1 | 0.002411 | 0.03208 | 9.5117 | 4.7621 | 0.0001 |
| Op.2 | 1 | 0.003243 | 0.03744 | 11.163 | 5.7723 | 0.0001 |
| Op.3 | 1 | 0.003394 | 0.04069 | 12.172 | 5.2423 | 0.0001 |
| Op.4 | 1 | 0.003090 | 0.03388 | 10.063 | 5.4314 | 0.0001 |

Df = Degrees of freedom, SS = Sum of squares, $F$ = $F$ statistics, $Z$ = Effect size

**Figure 4.**
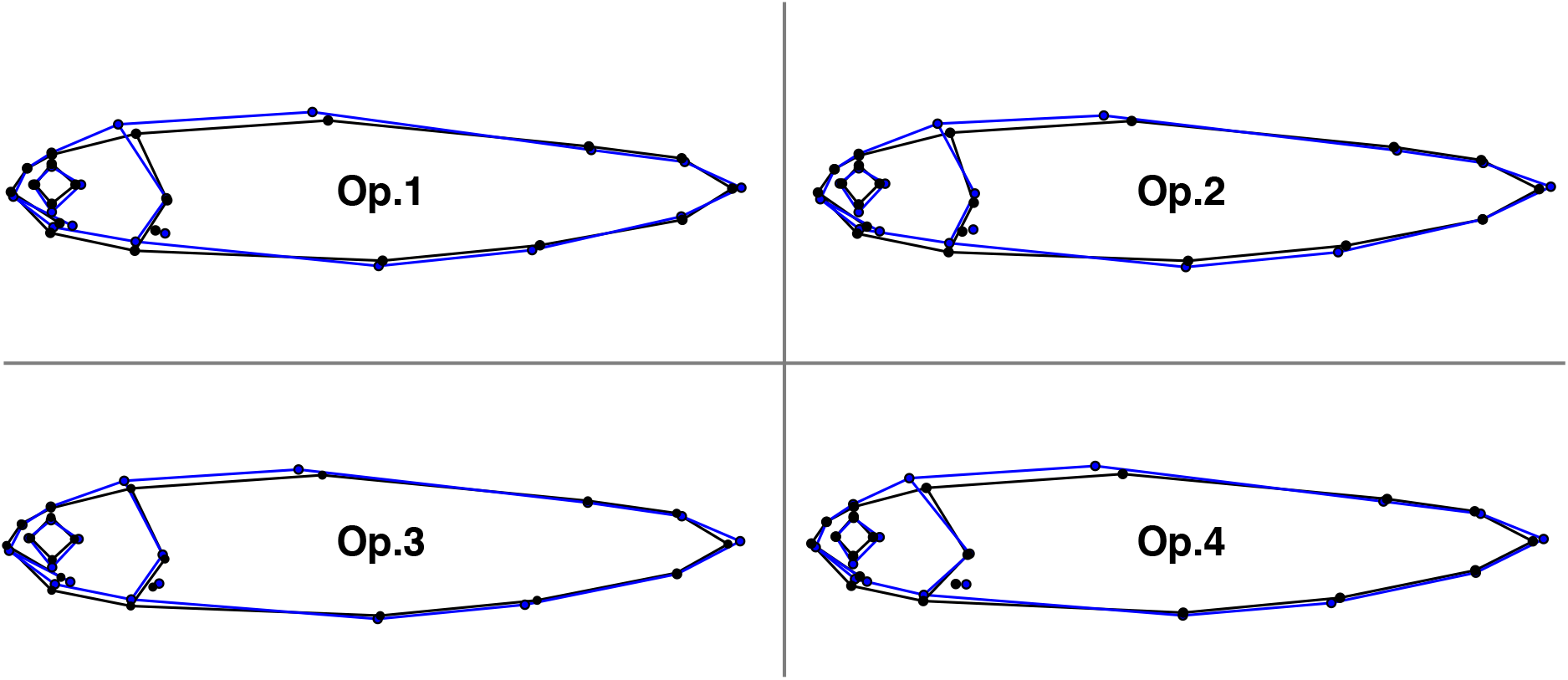
Comparisons of the mean body shape of 289 Atlantic salmon in the rivers Spey (black) and Oykel (blue) landmarked by four independent operators. Morphological differences were magnified six times to aid visualisation.

### Test 3. Were identified between-river differences consistent across all operators?

All comparisons of the “direction” of body shape variation between rivers were significant for all operators, meaning that the way in which shape differed between the rivers Spey and Oykel was approximately the same for all operators (Table 3). In contrast, pairwise comparisons of between-river and between-operator differences were mostly non-significant, with nine of 24 tests generating *P*-values < 0.05 (Supplementary Table 3). This indicates that the shape variation between rivers and operators were mainly divergent and not collinear.

**Table 3.** Pairwise tests of angles between body shape differences among rivers detected by four different operators. Measurements of angles (degrees) between bwgPC1 vectors (below the diagonal) and *P*-values (above the diagonal) are shown. Significant *P*-values (in bold) indicate that shape change vectors are similar to each other.

|  | Op.1<br>(Spey vs.<br>Oykel) | Op.2<br>(Spey vs.<br>Oykel) | Op.3<br>(Spey vs.<br>Oykel) | Op.4<br>(Spey vs.<br>Oykel) |
| --- | --- | --- | --- | --- |
| **Op.1**<br>(Spey vs. Oykel) |  | **5.3 x 10<sup>-20</sup>** | **1.8 x 10<sup>-14</sup>** | **6.0 x 10<sup>-17</sup>** |
| **Op.2**<br>(Spey vs. Oykel) | 22.7° |  | **1.1 x 10<sup>-22</sup>** | **1.9 x 10<sup>-19</sup>** |
| **Op.3**<br>(Spey vs. Oykel) | 28.4° | 17.0° |  | **1.9 x 10<sup>-19</sup>** |
| **Op.4**<br>(Spey vs. Oykel) | 24.3 ° | 20.8° | 20.8° |  |

Estimated mean distances between rivers computed through bootstrapping were similar across operators, as shown by the widely overlapping confidence intervals (Table 4), suggesting that different operators did not influence the magnitude of shape difference detected between the rivers Spey and Oykel.

**Table 4.** Estimated mean and median shape distance (with confidence intervals) between the rivers Spey and Oykel obtained by each operator.

| Operator | Mean distance | Median distance | Lower CI extreme | Upper CI extreme |
| --- | --- | --- | --- | --- |
| Op.1 | 0.00606 | 0.00608 | 0.00491 | 0.00729 |
| Op.2 | 0.00697 | 0.00695 | 0.00573 | 0.00821 |
| Op.3 | 0.00707 | 0.00706 | 0.00571 | 0.00845 |
| Op.4 | 0.00684 | 0.00684 | 0.00553 | 0.00821 |

### Test 4. Can divergent datasets from different operators be merged?

There were significant differences in body shape between fish from the rivers Spey and Oykel digitised separately by Op.2 and Op.4 (Table 5). Notably, shape variation explained by the rivers was markedly higher for these merged datasets compared to the between-river differences detected by single operators (*r*^2^ = 0.11-0.21 vs. 0.03-0.04, respectively; Tables 5 and 2). Similarly, the between-group PCA separated fish from different rivers with a higher accuracy than the analogous analysis performed on individual operator datasets (93.6% vs. 70.9% mean classification success rate, respectively; Supplementary Tables 4 and 2).

**Table 5.**
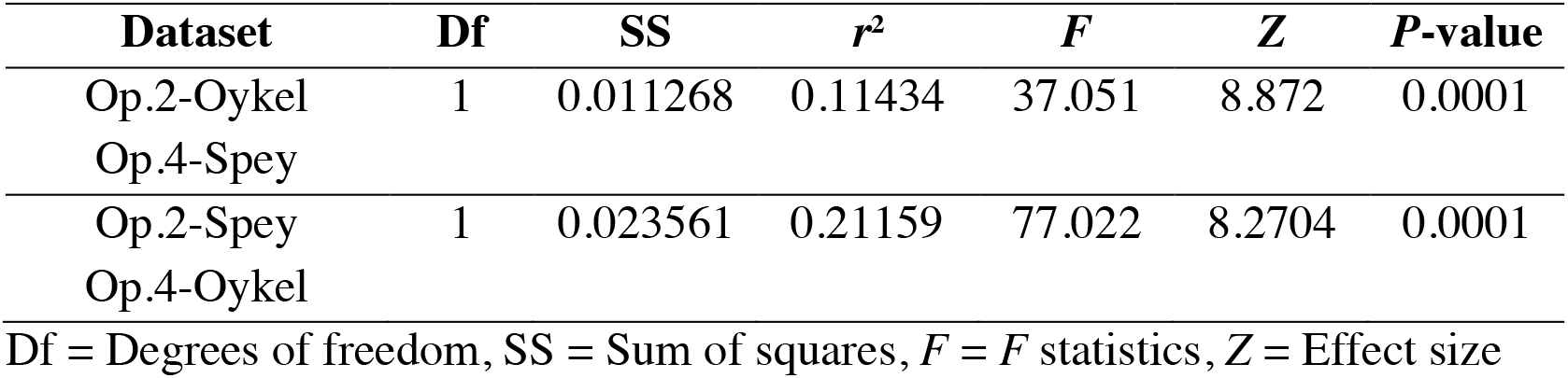
Procrustes ANOVA summary statistics of effect of river-of-origin on the body shape of 289 Atlantic salmon based on combined datasets of Op.2 and Op.4.

The comparison of the “direction” of between-river body shape variation detected by the two merged datasets from Op.2 and Op.4 was highly significant (*P*-value < 0.0001; Supplementary Table 5), meaning that the way in which shape differed between the rivers Spey and Oykel was approximately the same regardless of the selected dataset. However, only five of eight comparisons were found to be significant when comparing the two Op.2 and Op.4 merged datasets with the complete within-operator datasets including both rivers (Supplementary Table 5), indicating that the river differences detected by combined and individual operator datasets were only partly similar.

### Test 5. Quantifying intra-operator error

There was extensive overlap among landmarking trials, suggesting a high consistency in digitisation across all operators (Fig. 5). Pairwise comparisons supported this since none of the mean body shapes differed significantly between repeated digitisations (*P*-values > 0.82; Table 6). All four operators achieved the highest landmarking consistency between trials 2 and 3, as indicated by the smallest Euclidean distance values (0.001-0.003; Table 6).

**Table 6.** Pairwise comparisons of the body shape of 20 Atlantic salmon in three landmarking trials by four independent operators.

| Operator | Trials | Euclidean dist. | Hotelling's $T^2$ | $F$ | $P$-value |
| --- | --- | --- | --- | --- | --- |
| Op.1 | 1 vs. 2 | 0.00240 | 67.48440 | 0.1869 | 0.97 |
|  | 1 vs. 3 | 0.00315 | 141.9397 | 0.3932 | 0.87 |
|  | 2 vs. 3 | 0.00153 | 22.45716 | 0.0622 | 0.99 |
| Op.2 | 1 vs. 2 | 0.00358 | 47.28936 | 0.1310 | 0.99 |
|  | 1 vs. 3 | 0.00377 | 69.33009 | 0.1921 | 0.97 |
|  | 2 vs. 3 | 0.00236 | 56.09318 | 0.1554 | 0.98 |
| Op.3 | 1 vs. 2 | 0.00353 | 49.37887 | 0.1368 | 0.99 |
|  | 1 vs. 3 | 0.00360 | 50.734703 | 0.1405 | 0.98 |
|  | 2 vs. 3 | 0.00127 | 15.02887 | 0.0416 | 0.99 |
| Op.4 | 1 vs. 2 | 0.00922 | 184.9322 | 0.5123 | 0.82 |
|  | 1 vs. 3 | 0.00731 | 128.5921 | 0.3562 | 0.89 |
|  | 2 vs. 3 | 0.00344 | 58.91748 | 0.1632 | 0.98 |

**Figure 5.**
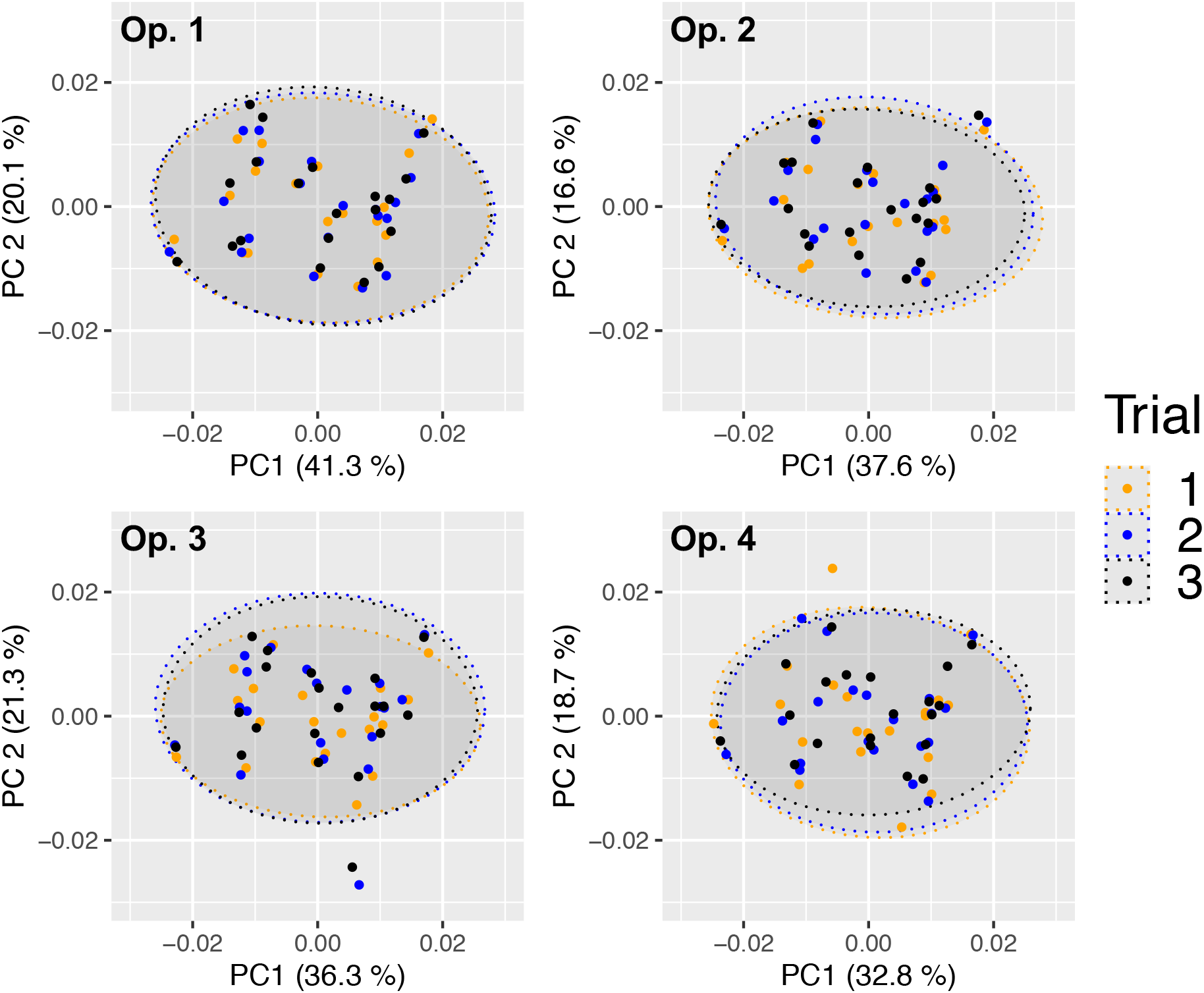
Principal components analysis scatterplots of Procrustes-aligned coordinates for 20 Atlantic salmon in three landmarking trials by four independent operators. Dots represent individual fish. Variance (%) explained by the first and second axes and 95% confidence ellipses are shown.

Repeatability was also high for all operators (0.925-0.977), indicating high landmarking precision (Table 7). Nested Procrustes ANOVA showed that 67.3% of the morphological variation within the subset of 20 fish was explained by individual fish (‘ID’), while 25.7% and 7.0% of the variation was attributable, respectively, to inter-(‘ID:operator’) and intra-(residual) operator digitisation error (Supplementary Table 6).

**Table 7.** Repeatability values for the three landmarking trials on 20 Atlantic salmon by four independent operators.

| Operator | Repeatability | Procrustes ANOVA $r^2$ (%) |
| --- | --- | --- |
| Op.1 | 0.977 | 95.3 |
| Op.2 | 0.961 | 92.4 |
| Op.3 | 0.956 | 91.5 |
| Op.4 | 0.925 | 86.3 |

## Discussion

We show here that independent operators applying an identical landmarking scheme to replicate photographs of live Atlantic salmon taken in field conditions yielded significantly different mean body shapes (*Test 1*). However, morphological differences between salmon from different rivers were detected by all operators (*Test 2*), and these were consistent differences across all operators (*Test 3*), provided they landmarked both rivers and not one each (*Test 4*). Furthermore, intra-operator error calculated on a subset of samples was minimal, suggesting that it did not have a significant influence on the body shape results obtained by the different operators (*Test 5*).

Despite digitising replicate photographs with homologous landmarks, all the operators produced significantly different mean body shapes. The high rate (83.0%) of specimens assigned to the correct operator by the between-group PCA suggests that the operators introduced a systematic error, which created four identifiable body shapes despite following the same landmarking scheme. This digitisation bias is likely to have been introduced by operators consistently applying personal, fine-scaled landmarking rules in addition to the general scheme. The fact that the differences among operators were localised mainly in the head region (landmarks 1-12) may be explained by the less discrete and recognizable nature of these landmarks compared to those located on well-defined anatomical loci, such as fin intersections (landmarks 13-20). This suggests that the use of unambiguous landmarks can be an effective way of reducing measurement error in GM (Fagertun et al., 2014; Campomanes-Álvarez et al., 2015, Fruciano et al., 2017).

In GM studies, digitisation is typically performed by a single operator, leaving the question of whether multiple operators digitising the same set of images would generate different results. This could undermine the reliability of findings presented by many GM investigations, particularly those using images of live animals taken in field conditions, which are potentially more prone to measurement error (Webster & Sheets, 2010; Muir, Vecsei & Krueger, 2012). In our study, however, we found that inconsistencies between operators did not mask small, but significant morphological differences between fish from the rivers Spey and Oykel, which were consistent across operators. The fact that, as shown by tests of angles and bootstrapped estimates of mean distances, all the operators detected analogous between-river differences, strongly suggests that they were biologically authentic. Similarly, Fruciano et al. (2020) found that preservation methods significantly affected the body shape of brown trout (*Salmo trutta* L.), but the subsequent between-groups classification was similar regardless of preservation method. As suggested by Fruciano et al. (2020), this could be because the shape variation detected by the operators between the rivers Spey and Oykel was not significantly affected by inter-operator differences in landmarking since they were not collinear (i.e. they followed different “directions”, as shown by the angle comparisons).

Conversely, merging landmark data of fish from the rivers Spey and Oykel digitised by two distinct operators (Op.2 and Op.4, *Test 4*) had a significant impact on subsequent analyses and produced contrasting results. As shown by Procrustes ANOVA and between-group PCA classification rate, shape differences between rivers in the merged datasets were greater than those detected by single operators, suggesting they were artificially inflated by inter-operator digitisation error. Angle comparisons showed that the river differences detected by combined and individual operator datasets were partly dissimilar. Overall, these findings point towards a potential risk in pooling datasets from multiple operators when there are confounding biological factors, as highlighted by other studies (Fruciano et al., 2017; Evin, Bonhomme & Claude, 2020). Distinct operators obtained analogous results when they landmarked both rivers (and not one river each as in *Test 4*). This suggests that operators digitising at least a sub-set of all data groups of interest (rivers in this case) may be an effective way of mitigating inter-operator error and potentially enabling data sharing.

In contrast to the inter-operator effects described in this study, we found no statistical evidence of intra-operator effects on the quantification of fish morphology. On the contrary, we found a very high level of repeatability across trials for all operators. This corroborates previous studies that showed intra-operator error to be limited (e.g. Cardoso & Saunders, 2008; Dujardin, Kaba & Henry, 2010; Wilson, Cardoso & Humphrey, 2011; Robinson & Terhune, 2017; Shearer et al., 2017; Thoma et al., 2018; Fox, Veneracion & Blois, 2020). Interestingly, for all operators, landmarking consistency was highest between their last two trials, suggesting that they ‘learnt’ where to place the landmarks with increasing experience of the images. However, it should be noted that the first trial was performed while digitising all specimens, whereas trials 2 and 3 were performed consecutively after digitising the full dataset, which may have artificially inflated precision, with operators “remembering” their landmarking choices in trial 2 during trial 3.

The negligible impact of intra-compared to inter-operator error was also clearly shown by the percentage of variance explaining shape variation in the sub-sample of 20 fish (*Test 5*, 7.0% vs. 25.7%, respectively). Interestingly, the percentage of variance explained by inter-operator error (25.7%) is similar to that reported by Fruciano et al. (2020) for brown trout photographed in the field (30.1%), and supports previous studies that identified inter-operator effects as the major source of error in GM analyses (Ross & Williams, 2008; Dujardin, Kaba & Henry, 2010; Campomanes-Álvarez et al., 2015; Fruciano, 2016; Shearer et al., 2017; Fox, Veneracion & Blois, 2020).

## Conclusions

Overall, we show that, even when significant inter-operator error is introduced through digitisation, following an identical landmarking scheme can be an effective tool to obtain robust and reliable results, even without accounting for variation introduced by the photography process, which was not quantified here. This implies that GM studies based on common landmarking schemes are potentially reproducible, even when analyses are based on images of live specimens taken in the field, as in the current study. Nevertheless, since operator error can vary between studies and is impossible to determine *a priori*, we recommend assessing the magnitude and effects of landmarking error by using multiple operators for a sub-set of samples, as here, to improve confidence in study results. If landmark data merging is required, we recommend that all the operators involved digitise at least a sub-set of all data groups of interest (rivers in this case) to mitigate inter-operator error.

## Acknowledgements

We would like to thank the Spey Fisheries Board and the Kyle of Sutherland Fisheries Trust for their help and support with this project. We would also like to thank Jonathan Archer, Georgios Kyriakou, Fraser Brydon, Jessica Whitney and Mustafa Soganci for their help collecting the data for this study. We also thank Carmelo Fruciano, Michelle C. Gilbert and Kiran Liversage for their constructive reviews.

