## Supplementary material for "Is shape in the eye of the beholder? Assessing landmarking error in geometric morphometric analyses on live fish"

**Supplementary Table 1.** Operator classification obtained by Between-group PCA using the leave-one-out cross-validation. Numbers represent fish specimens assigned to operators based on their morphology. Percentage of classification accuracy is shown.

| | **Op.1** | **Op.2** | **Op.3** | **Op.4** | **Overall classification accuracy** |
|---|---|---|---|---|---|
| **Op.1** | 222 | 16 | 30 | 21 | 76.8 % |
| **Op.2** | 15 | 267 | 3 | 4 | 92.4 % |
| **Op.3** | 35 | 8 | 220 | 26 | 76.1 % |
| **Op.4** | 19 | 4 | 16 | 250 | 86.5 % |

**Supplementary Table 2.** River classification obtained by Between-group PCA for each operator using the leave-one-out cross-validation. Numbers represent specimens assigned to rivers based on their morphology. Percentage of classification accuracy is shown.

| **Operator** | **River of origin** | **Assigned to Oykel** | **Assigned to Spey** | **Overall classification accuracy** |
|---|---|---|---|---|
| Op.1 | Oykel | 113 | 32 | 73.4% |
| | Spey | 45 | 99 | |
| Op.2 | Oykel | 106 | 39 | 70.9% |
| | Spey | 45 | 99 | |
| Op.3 | Oykel | 103 | 42 | 70.6% |
| | Spey | 43 | 101 | |
| Op.4 | Oykel | 97 | 48 | 68.5% |
| | Spey | 43 | 101 | |

**Supplementary Table 3.** Pairwise tests of angles between body shape differences among rivers (Spey vs. Oykel) and operators (e.g., mean shape Op.1 - mean shape Op.2). Significant *P*-values (in bold) indicate that shape change vectors are similar to each other.

| Comparison | | ***P*-value** | **Angle (degrees)** |
|---|---|---|---|
| Op.1 (Spey vs. Oykel) | Op.1 - Op.2 (mean shapes) | 0.227 | 83.1 |
| Op.1 (Spey vs. Oykel) | Op.1 - Op.3 (mean shapes) | **0.026** | 72.2 |
| Op.1 (Spey vs. Oykel) | Op.1 - Op.4 (mean shapes) | **0.021** | 71.4 |
| Op.1 (Spey vs. Oykel) | Op.2 - Op.3 (mean shapes) | **0.034** | 73.3 |
| Op.1 (Spey vs. Oykel) | Op.2 - Op.4 (mean shapes) | **0.034** | 73.3 |
| Op.1 (Spey vs. Oykel) | Op.3 - Op.4 (mean shapes) | 0.193 | 81.2 |
| Op.2 (Spey vs. Oykel) | Op.1 - Op.2 (mean shapes) | 0.425 | 88.3 |
| Op.2 (Spey vs. Oykel) | Op.1 - Op.3 (mean shapes) | **0.048** | 74.7 |
| Op.2 (Spey vs. Oykel) | Op.1 - Op.4 (mean shapes) | **0.035** | 73.3 |
| Op.2 (Spey vs. Oykel) | Op.2 - Op.3 (mean shapes) | 0.188 | 81.8 |
| Op.2 (Spey vs. Oykel) | Op.2 - Op.4 (mean shapes) | 0.128 | 79.5 |
| Op.2 (Spey vs. Oykel) | Op.3 - Op.4 (mean shapes) | 0.205 | 82.4 |
| Op.3 (Spey vs. Oykel) | Op.1 - Op.2 (mean shapes) | 0.345 | 86.3 |
| Op.3 (Spey vs. Oykel) | Op.1 - Op.3 (mean shapes) | 0.083 | 77.2 |
| Op.3 (Spey vs. Oykel) | Op.1 - Op.4 (mean shapes) | 0.110 | 78.7 |
| Op.3 (Spey vs. Oykel) | Op.2 - Op.3 (mean shapes) | 0.291 | 84.9 |
| Op.3 (Spey vs. Oykel) | Op.2 - Op.4 (mean shapes) | 0.267 | 84.3 |
| Op.3 (Spey vs. Oykel) | Op.3 - Op.4 (mean shapes) | 0.351 | 86.5 |
| Op.4 (Spey vs. Oykel) | Op.1 - Op.2 (mean shapes) | 0.345 | 86.3 |
| Op.4 (Spey vs. Oykel) | Op.1 - Op.3 (mean shapes) | **0.049** | 74.8 |
| Op.4 (Spey vs. Oykel) | Op.1 - Op.4 (mean shapes) | **0.017** | 70.5 |

| | | | |
|---|---|---|---|
| Op.4 (Spey vs. Oykel) | Op.2 - Op.3 (mean shapes) | 0.088 | 77.5 |
| Op.4 (Spey vs. Oykel) | Op.2 - Op.4 (mean shapes) | **0.046** | 74.6 |
| Op.4 (Spey vs. Oykel) | Op.3 - Op.4 (mean shapes) | 0.124 | 79.3 |

**Supplementary Table 4.** River classification obtained by Between-group PCA for merged datasets of operators 2-4 using the leave-one-out cross-validation. Numbers represent specimens assigned to rivers based on their morphology. Percentage of classification accuracy is shown.

| **Operator** | **River of origin** | **Assigned to Oykel** | **Assigned to Spey** | **Overall classification accuracy** |
|---|---|---|---|---|
| Op.2 | Oykel | 133 | 12 | 92.7% |
| Op.4 | Spey | 9 | 135 | |
| Op.4 | Oykel | 132 | 13 | 94.5% |
| Op.2 | Spey | 3 | 141 | |

**Supplementary Table 5.** Pairwise tests of angles between body shape differences among rivers detected by merged datasets of Ops. 2-4 as well as by the separate datasets of all four operators. Measurements of angles (degrees) between bwgPC1 vectors (below the diagonal) and *P*-values (above the diagonal) are shown. Significant *P*-values (in bold) indicate that shape change vectors are similar to each other.

| | Op.2-Oykel Op.4-Spey | Op.2-Spey Op.4-Oykel | Op.1 (Oykel vs. Spey) | Op.2 (Oykel vs. Spey) | Op.3 (Oykel vs. Spey) | Op.4 (Oykel vs. Spey) |
|---|---|---|---|---|---|---|
| Op.2-Oykel Op.4-Spey | | **$1.4 \times 10^{-6}$** | 0.445 | 0.093 | **0.022** | 0.304 |
| Op.2-Spey Op.4-Oykel | 48.7° | | **$3.0 \times 10^{-7}$** | **$2.9 \times 10^{-5}$** | **0.001** | **$4.4 \times 10^{-7}$** |
| Op.1 (Oykel vs. Spey) | 88.7° | 46.4° | | **$5.3 \times 10^{-20}$** | **$1.8 \times 10^{-14}$** | **$6.0 \times 10^{-17}$** |
| Op.2 (Oykel vs. Spey) | 77.8° | 54.2° | 22.7° | | **$1.1 \times 10^{-22}$** | **$1.9 \times 10^{-19}$** |
| Op.3 (Oykel vs. Spey) | 71.5° | 61.0° | 28.4° | 17.0° | | **$1.9 \times 10^{-19}$** |
| Op.4 (Oykel vs. Spey) | 85.3° | 46.9° | 24.3 ° | 20.8° | 20.8° | |

**Supplementary Table 6.** Nested Procrustes ANOVA summary statistics of effect of fish specimen ('ID') and operators on shape variation after three repeated digitisations of twenty salmon smolts (Df = Degrees of freedom, SS = Sum of squares, *F* = *F* statistics, *Z* = Effect size).

| | **Df** | **SS** | **$r^2$** | ***F*** | ***Z*** | ***P*-value** |
|---|---|---|---|---|---|---|
| ID | 19 | 0.062678 | 0.67295 | 80.8462 | 30.227 | 0.0001 |
| ID:Operator | 60 | 0.023932 | 0.25695 | 9.7753 | 14.920 | 0.0001 |
| Residuals | 160 | 0.006529 | 0.07010 | | | |
| Total | 239 | 0.093139 | | | | |

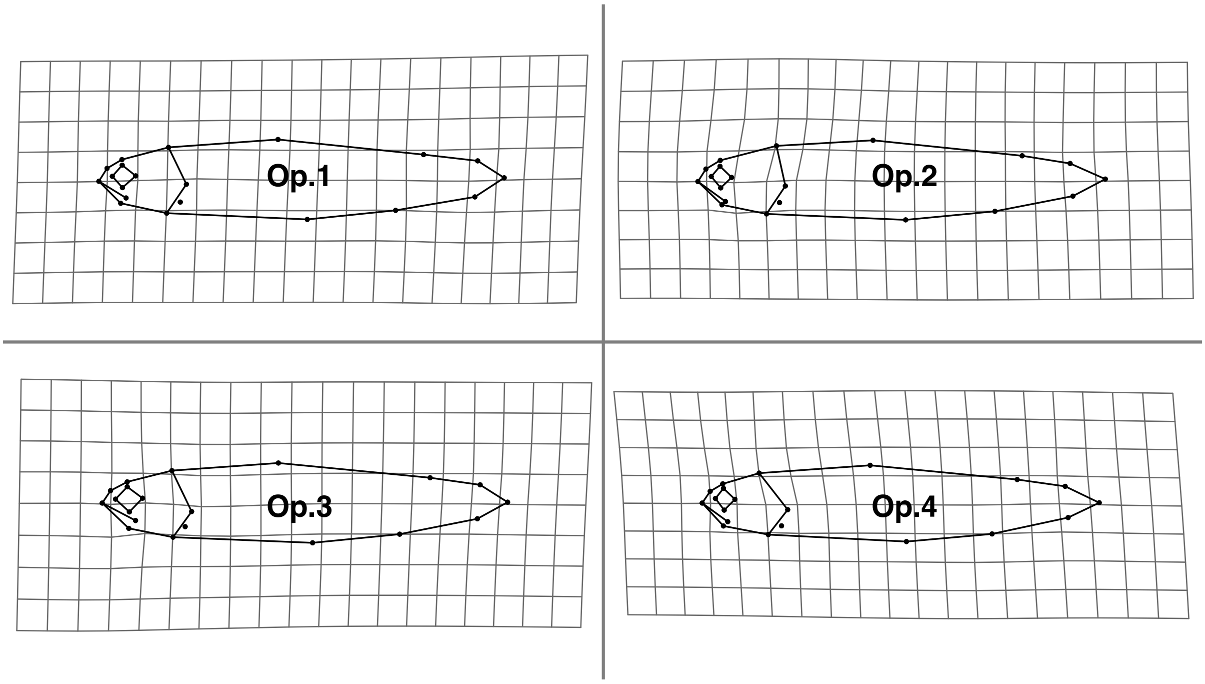


**Supplementary Figure 1.** Differences in shape between operators illustrated using thin-plate spline deformation grids. Mean body shapes for each operator are projected against the overall shape calculated for all specimens from all four operators. Morphological differences were magnified two times for easier visualisation.

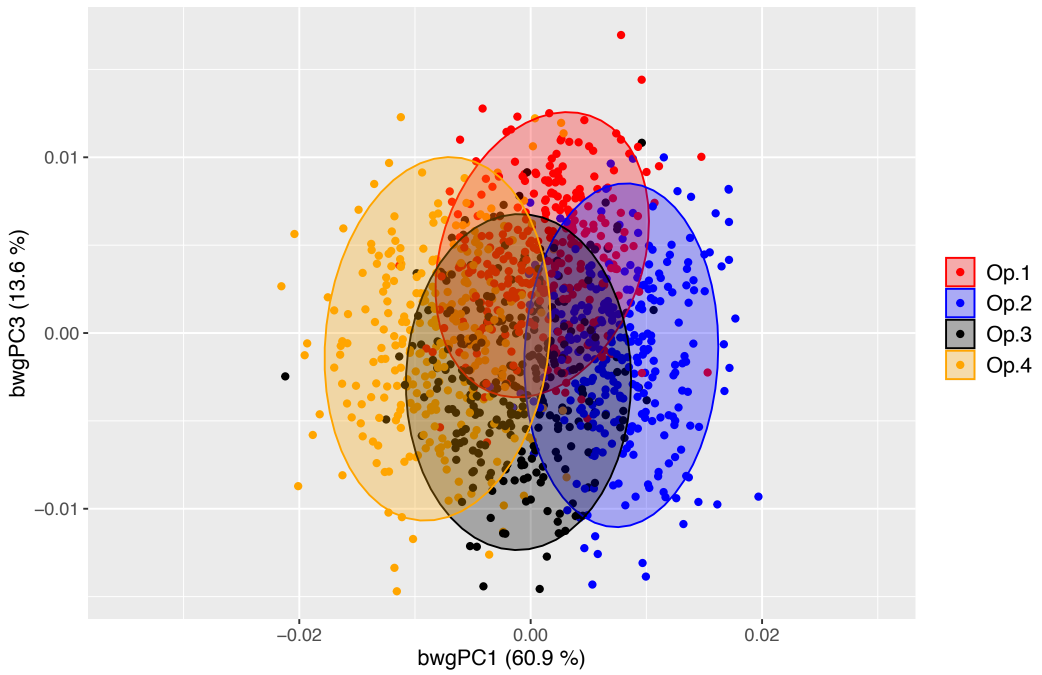


**Supplementary Figure 2.** Between-operator PCA scatterplot showing the cross-validated scores along the first and third between-group principal components (bwgPCs). Dots represent individual Atlantic salmon ($n$ = 289) landmarked by four independent operators (different colours). Between-operator variance (%) explained by the first and third axes is shown.

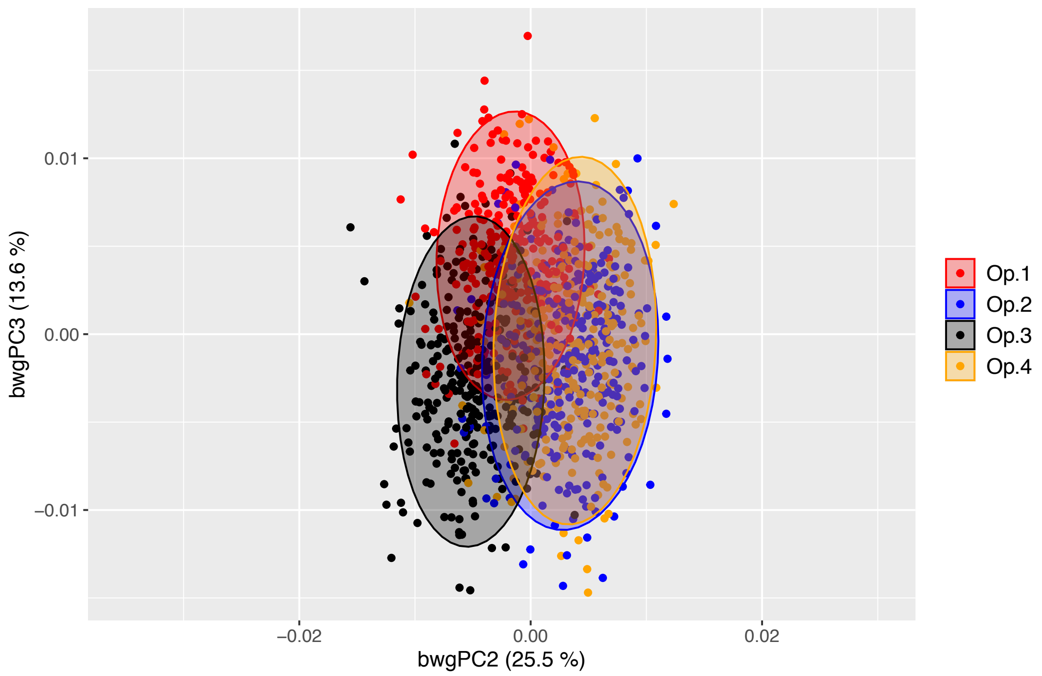


**Supplementary Figure 3.** Between-operator PCA scatterplot showing the cross-validated scores along the second and third between-group principal components (bwgPCs). Dots represent individual Atlantic salmon ($n$ = 289) landmarked by four independent operators (different colours). Between-operator variance (%) explained by the second and third axes is shown.
